# Whole genome linkage analysis in a large Brazilian multigenerational family reveals distinct linkage signals for Bipolar Disorder and Depression

**DOI:** 10.1101/106260

**Authors:** Mateus Jose Abdalla Diniz, Andiara Calado Saloma Rodrigues, Ary Gadelha, Shaza Issam Alsabban, Camila Guindalini, Jose Paya-Cano, Simone de Jong, Peter McGuffin, Rodrigo A. Bressan, Gerome Breen

## Abstract

Both common and rare genetic variation play a role in the causes for mood disorders. Very large families pose unique opportunities and analytical challenges but may provide a way to identify regions and mutations associated with mood disorders. We identified a family with a high prevalence (~30%) of mood disorders in a rural village in Brazil, featuring decreasing age of onset over generations. The pattern of inheritance was complex with 32 Bipolar type I cases, 11 Bipolar type II and 59 recurrent and/or severe Depression cases in addition to other phenotypes. We enrolled 333 participants with DNA samples from a broader pedigree of 960 subjects for genotyping using the Affymetrix 10K array. Non-parametric linkage was carried out via MERLIN and parametric with both MERLIN and MCLINKAGE. We exome sequenced a subset of the family (n=27) in order to identify rare variation within the linkage regions shared by affected family members. We identified four genome wide significant and four suggestive linkage regions on chromosomes 1, 2, 3, 11 and 12 for different phenotype definitions. However, no region received strong joint support in both the parametric and non-parametric analyses. Exome sequencing revealed potential deleterious variants in 11p15.4 for MDD and 1q21.1-1q21.3 and 12p23.1-p22.3, implicated in cell signaling, adhesion, translation and neurogenesis processes. Overall, our results suggest promising, but not definitive or confirmed evidence, that rare genetic variation contributes to the high prevalence of mood disorders in this multi-generational family. We note that a substantial role for common genetic variation is likely given the strength of the linkage signals observed.

The World Health Organisation reports depression and bipolar disorder as the second and seventh most important causes of years lost due to disability worldwide[1]. The heritability of bipolar disorder is between 60-90% with a lower but still substantial heritability for major depression (40-45%) [2]; [3]. First-degree relatives of bipolar disorder probands have a 5-10 fold increase in risk of developing the illness compared to relatives of controls but also show a three fold increase in unipolar depression, indicating that bipolar disorder does not “breed true” [4]. Large collaborative genome-wide association studies (GWAS) have uncovered several common genetic variants of small effect [5]. Genomewide estimates of heritability suggest that up to 60% of the genetic risk is contributed by common variants [6]. Overall, the current picture for bipolar disorder (and almost all complex traits) is a genetic architecture formed of both common and rare variants.

Linkage studies have been pursued on the basis that there may be variants of greater effect shared between and within affected families. However these studies have usually focused on collections of comparatively small families or sib pairs and few consistent findings have emerged [7]. Large multigenerational families (e. g. of >30 affected individuals) theoretically offer a powerful means for mapping complex disease loci that are individually rare but common in a single family. These loci may be more highly penetrant and of larger effect than loci found with GWAS [8]. Here we report the results of the Brazilian Bipolar Family (BBF) study on a five-generation family of 639 members of which 333 were enrolled in the current analyses. Our objectives were to perform a linkage analysis with genome coverage and try to identify new genes/mutations related to bipolar and other mood disorders in the family. Here we report our findings and preliminary results of sequencing of linkage regions.

## Methods

### Family Ascertainment

The Brazilian Bipolar Family (BBF) consists of 960 members. Ascertainment was via a 45-year-old female proband with severe Type-1 BP, who was treated by one of the psychiatrists involved in the study (M. D). She stated that there were dozens of cases of mood disorders in the family, most of whom lived in a small village in a rural area of a state north of São Paulo. Cooperation from the family and a within family published book about the history of the family, self-published within the family, was invaluable for our ascertainment.

The grandparents of the proband were reported to be first cousins and both suffered from BP1 disorder. Of their 13 children, 12 had a confirmed bipolar mood disorder and many of them went on to have affected children. Notably, one of them went on to give birth to 14 affected children. The BBF also exhibits other features, including anticipation; an apparent pattern of earlier age of onset in affected individuals in successive generations.

Family members >16 years of age underwent semi-structured interviews, using the Portuguese version of the Structured Clinical Interview for DSM-IV Axis I Disorders (SCID) [9]. Members aged 6-16 were assessed using the Portuguese version of Kiddie-SADS-Present and Lifetime Version (K-SADS-PL) [10]. In total 308 interviews were completed, and 5 eligible members declined an interview. Discrepancies in diagnoses were reviewed by two independent psychiatrists and a final consensus diagnosis was assigned. The family had an obvious division, with densely affected large branches from the village and nearby forming a core that we termed “Branch 1” and the rest of the family, which has less densely affect branches, and had migrated to urban areas around south-eastern Brazil.

### Phenotype Models

Three phenotypic models were constructed for the analyses: a narrow, broad, and super model, in addition to a depression only model. The narrow affection model included family members that fulfilled DSM-IV criteria for BPI, BPII, or schizoaffective disorder. The broad model included family members in the narrow model in addition to family members who fulfilled DSM-IV criteria for BPNOS and cyclothymia. The super model included cases from the broad model and those with one or more episodes of major depression of moderate to severe MDD or who fulfilled diagnostic criteria for dysthymia as defined by DSM-IV. Finally, family members were included in the depression model if they had a history of dysthymia or experienced one episode of major depression (Table 1). Non interviewed family members were given the status “unknown”. Individuals who were interviewed and did not receive any mood (or other psychiatric) disorder diagnosis were labelled as “unaffected” in the analyses.

**Table 1.**
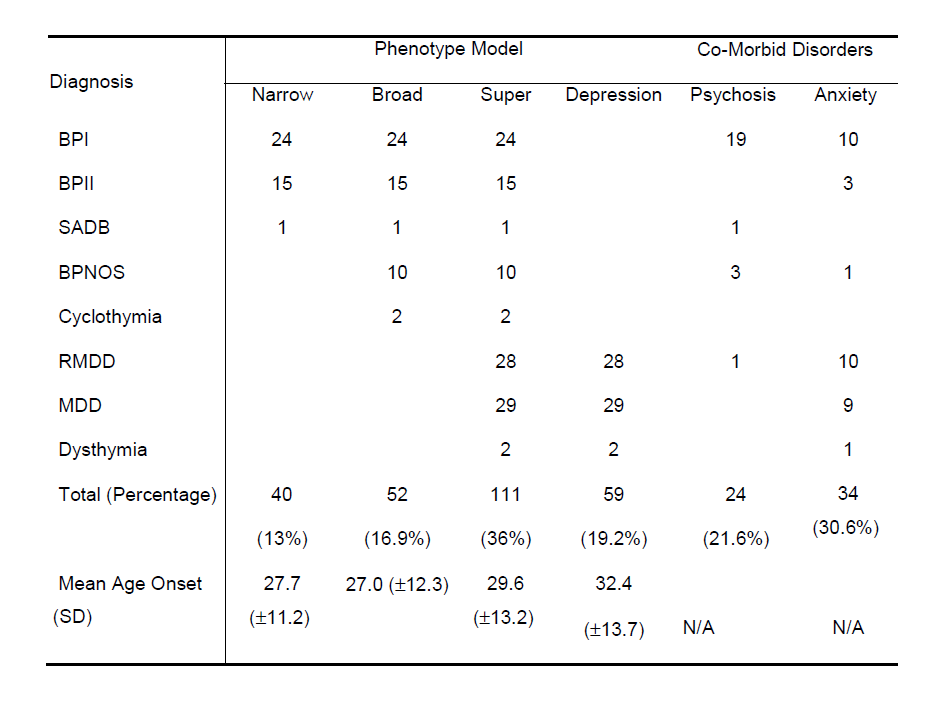
**Phenotype models by number and percentage of affected family members and co-morbid psychosis and anxiety disorders.** The total number of family members affected with a mood disorder is 111. The total number of interviews conducted is 308.

### Sample Collection and Genotyping

Following diagnostic interview, interviewers obtained 30ml of whole blood in 4 7.5 ml (EDTA containing) monovettes for adults and lesser amounts or saliva for 8 adults and 17 children given personal preference or age (DNA Genotek Inc., Ontario, Canada). Genomic DNA was isolated from whole blood and saliva at the Federal University of São Paulo using standard procedures. In total, 333 DNA samples were extracted, of which 324 were genotyped using the Affymetrix 10K microarray. One sample with discordant sex information was removed from the data set.

We excluded SNPs on genotype missingness (>10%), a minor allele frequency <25% in BBF founders (n=54) and deviation from HWE in founders only (p<10^−4^). 5315 SNPs remained available for analysis. Pedigree structure, Mendelian and non-Mendelian errors were estimated using PLINK pair-wise IBD estimation, PEDSTATS [11] and MERLIN [12] with MERLIN’s “pedwipe” function used to remove them. We also used the McLinkage Check Errors procedure to calculate the posterior probability of genotype mistyping at each marker in the data (given the observed genotypes in relatives), which were incorporated into subsequent McLinkage analyses.

The BBF members self reported mixed Southern European ancestry, which was confirmed by the interviewers impressions. Genetic analysis of principal components, as implemented in EIGENSOFT version 3.0 (http://www.hsph.harvard.edu/alkes-price/software/) revealed that family members clustered closely with the Northern and Western European and Tuscan Italian populations (data not shown).

### Structure of Analysis

Following completion of the QC procedures, 301 BBF members and 5315 were available for analysis. Four clean pedigree files were generated for the analyses: (a) the Branch 1 (village) sub-families, (b) All BBF sub-families, (c) Branch 1 with structure intact and (d) Total BBF with structure intact.

### Inheritance Models

Guided by large family studies from similar populations [13],[14], we specified dominant and recessive models. We assumed 1% penetrance for zero copies of the disease allele (i.e. phenocopy rate), 81% for one copy, and 90% for two copies, and 1%,1% and 90% for the recessive model. Under the dominant model, disease allele frequencies assumed were 0.003, 0.03, and 0.13 for the narrow, broad, and super phenotype models (reflecting prevalence estimates of 1.5%, 5% and 20%). Under the recessive model, disease allele frequencies assumed were 0.07, 0.03, and 0.46 for each phenotype.

We also performed analyses on a depression only model. Using a population prevalence for recurrent or long (>6 months) single episode severe major depression of 5%, and a penetrance estimate of 50%, we specified disease allele frequency of 0.005 and parameter penetrances of 5%, 50% and 50% for a dominant model and 5%, 5% and 50% for a recessive model with a disease allele frequency of 0.05 (dominant) and 0.33 (recessive) as previously used for the analysis of depression pedigrees by our group [2]. Genotypes of BBF founders allowed estimation of population specific marker allele frequencies [15]. We use the map provided with the array, with additional checking and mapping of SNPs to the reference genome (data not shown).

### Parametric Linkage Analysis using McLinkage

Two-point parametric linkage on the BBF and Branch 1 was conducted using TwoPointLods, part of the McLinkage package. For reasons of computation, regions reaching suggestive or significant thresholds in 2-point linkage were analysed using McLinkage multipoint parametric linkage, which samples from the posterior distribution of inheritance vectors, given the observed genotypes, and calculates multipoint LOD scores (MLOD) and TLOD scores between the specified phenotype and genetic markers in a pedigree [16].

### Merlin Analyses

Linkage analysis programs, such as MERLIN, that use the Lander-Green [17] exact LOD score calculation require large pedigrees to be split. We first split the BBF into 19 (<30 bit) sub-families, preserving family relatedness, such as first cousin marriages and complex marriage loops where possible, prior to breaking these loops. Two-point and multipoint parametric linkage analysis was conducted on 19 subfamilies belonging to all three branches of the family, and 12 subfamilies belonging to Branch 1 of the family using the models described above. Heterogeneity LOD (HLOD) scores were also calculated.

We also employed MERLIN to conduct multipoint NPL methods of linkage analysis, examining both affected relative pairs, NPL_pairs_, and affected subgroups within the family, NPL_all_, for allele sharing by descent. We selected the Kong and Cox (1997) exponential model flag in MERLIN (–exp) to convert NPL_pairs_ and NPL_all_ Z-scores to LOD scores and used MERLIN –trim to drop non-informative family members from the analysis.

### Whole-genome Linkage Thresholds

We used a threshold of LOD ≥ 3 and LOD ≥ 2 as indicative of whole-genome significant and suggestive linkage in our analyses [18]. We corrected for multiple testing using the formula of Ott, as used in [2]

LOD − log_10_(number of independent tests performed)

We used the Matrix Spectral Decomposition (matSpD) method to estimate the equivalent number of independent tests performed in our analyses (http://gump.qimr.edu.au/general/daleN/matSpD/) via use a correlation matrix of the LOD scores ≥ 2 achieved in all analyses to reflect a balanced liberal-conservative approach. This gave the independent tests performed as 8 and 5 for the parametric and NPL analyses respectively. Thus, corrected LOD scores of 3.90 and 2.90 are used to indicate significant and suggestive linkage in our parametric analyses and a corrected LOD score of 3.70 and 2.70 to indicate significant and suggestive linkage in our NPL analyses. Within the MERLIN results alone, very similar/near-identical estimates were generated via 100 simulations of the family in MERLIN preserving the structure of the pedigree, prior to splitting the sub-pedigrees before re-performing all NPL and parametric analysis. For reasons of computation, we were unable to do this confirmatory analysis in MCLINKAGE.

### Sequencing Methods

Paired-end 100bp whole exome sequencing on Illumina HiSeq 2000 for 29 individuals was outsourced to BGI (Beijing Genomics Institute). In brief, SOAPaligner was used for alignment to hg19 using default settings and allowing a maximum of 3 mismatches and SOAPsnp for assembly of consensus sequence and genotype calls, using default prior probabilities (novel hom 0.0005 and novel het 0.001) and refining SNP calling using known information (-2 option). Upon reception, files were converted to VCF4.0 and additional filters were applied: minimum read depth 30, maximum read depth 300 and quality 30 or higher using VCFtools v0.1.12a [19]. Ts/Tv ratios of remaining variants fell between 2.2 and 2.4. 2 individuals were excluded due to overall poor QC metrics. SNP and indels within linkage regions were then annotated using ANNOVAR [20] May 2014 version and databases.

Variants were filtered using the following criteria; <1% or <0.1% in any population in the Exome Aggregation Database and functional relevance (non-synonymous exonic SNV, stopgain or stoploss for SNVs). Finally, we required the variant to be present in at least 2 affected individuals for the particular phenotype and in a maximum of 1 unaffected and 1 married-in individual. In addition, we filtered our variants against a Brazilian population dataset, consisting of 604 exomes available from collaborators (see acknowledgments). We also screened the mutations in the EXAC database (http://exac.broadinstitute.org/ 11/08/15, see acknowledgments), which includes a large number of psychiatric exomes.

## Results

### Socio-demographics and Psychiatry Phenotype description

In total 111 (36%) interviewed family members received a mood disorder diagnosis. Forty (13%) family members fulfilled criteria for BPI, BPII, or SAD, which increased to 52 (16.9%) when BPNOS and cyclothymia were considered. More women (62.9%) reported mood disorders and BPI and BPII (61.3%). Unipolar depression was 17.3%; primary anxiety disorder prevalence was 13.2%; and alcohol abuse was 20.3% but not associated with mood disorders (data not shown).

One family member had schizoaffective disorder, four family members had learning disability, six had adjustment disorder, and five children were diagnosed with Attention Deficient Hyperactivity Disorder. The BBF also had higher than expected rates of autoimmune hypothyroidism (7.8%), type I diabetes (10.1%) and Parkinson’s disease (2.6%).

### Summary of Whole-Genome Linkage and Sequencing Results

Using the corrected whole-genome significant and suggestive thresholds, four chromosomal regions resulted in whole-genome significant LOD scores: 2p23.1-p22.3, 3p25.3-p24.1, 11p15.4, and 12q24.22-q24.32 and four chromosomal regions resulted in whole-genome suggestive LOD scores: 1p22.2-p21.3, 1q21.1-q21.3, 12p13.32-p13.31, and 22q11.21-q12.1 (Table 2). MERLIN multipoint parametric and non-parametric analyses generally did not yield similar results, although some support across linkage methodologies was observed for the regions on chromosome 11p15.4 and 22q11.21-q12.1 with LOD scores greater or equal to 1.0 for both approaches. In addition, most of the positive linkage results were derived from the Branch 1 subfamilies. Linkage peaks either remained the same or decreased in size with the inclusion of subfamilies from Branches 2 or 3 in the BBF analyses. The exception is chromosome 1q21.1-q21.3, where a suggestive linkage peak (maximum LOD=2.83) was only achieved in analysis of the BBF subfamilies.

**Table 2.** **Linkage regions identified.** Regions included in this table are those with suggestive or significant genome-wide linkage in any analytic configuration. Two asterisks denote SNPs with whole-genome significance and one asterisk denotes SNPs with whole-genome suggestive evidence for linkage. All of the maximum LOD scores presented in the table are from analyses conducted on Branch 1, except the maximum LOD score for chromosomal region 1q21.1-q21.3 is from analysis conducted on the BBF.

| Phenotype Model | Number Affected | Chromosomal Region | Software | Test | Mode Transmission | Maximum LOD |
| --- | --- | --- | --- | --- | --- | --- |
| Narrow | 40 | 1p22.2-p21.2 | MERLIN | NPL<sub>all</sub> | N/A | 2.96* |
|  |  | 3p24.3-p24.1 | McLinkage | TLOD | Dominant | 4.18** |
| Broad | 52 | 1p22.2-p21.3 | MERLIN | NPL<sub>all</sub> | N/A | 2.93* |
|  |  | 22q11.21-q12.1 | MERLIN | HLOD | Recessive | 3.76* |
| Super | 111 | 1q21.1-q21.3 | MERLIN | NPL<sub>pairs</sub> | N/A | 2.83* |
|  |  | 2p23.1-p22.3 | MERLIN | NPL<sub>pairs</sub> | N/A | 3.83** |
| Depression | 59 | 11p15.4 | MERLIN | NPL<sub>all</sub> | N/A | 4.49** |
|  |  | 12p13.32-p13.31 | McLinkage | TLOD | Dominant | 3.09* |
|  |  | 2q24.22-q24.32 | McLinkage | TLOD | Dominant | 4.74** |

Our sequencing of these regions is outlined in Table 3 with counts of variants that passed our filtering criteria per linkage region. We report only the variants with a GenomicSuperDup score <0.9. We required a minimum of 2 cases to carry each mutation. For the narrow phenotype no variants passed our sequencing filters. Notably we had a smaller number of cases sequenced for this phenotype (6 narrow cases versus 14 cases for depression). For the broad phenotype, no variants passed our sequencing filters for the broad phenotype. Linkage analysis had revealed two suggestive and one significant linkage peak for depression and the locus on chr11p15.4 was found to contain 3 predicted damaging non-synonymous variants in *ART5 and DCHS1*. One variant in the chr12q24.22-12q24.32 locus passed our filtering criteria in the *TCTN2* gene. For the super phenotype, the linkage region on chromosome 1q contains many segmental duplications, but after filtering for this, we still find some genes. We find three different variants predicted to be damaging in *ITGA10*, *ANP32E*, and in *TCHH*. On chromosome 2p23.1-p22.3, we found a damaging non-synonymous mutation in *FAM98A*. Unusually in EXAC (exac.broadinstitute.org), the position has two more alternate alleles observed in a few individuals. This meant all 4 bases were represented at this site, albeit 3 of them at very low allele frequency (aggregate 3/121378, i.e. about one in 40,000). The site has very good sequence coverage in our data and in EXAC. Examination of EXAC data Integrative Genome Viewer plots showed that these variants had good quality. EXAC has a density of 1 variant per 6bp and 7% of observed variants are multiallelic. (D. MacArthur personal communication, plots available upon request)

**Table 3.** **Sequencing results from exome sequencing of family members**. The first and second column contain the linkage region and the relevant phenotype. The variant location is stated in column three and four. All variants were annotated using ANNOVAR (http://annovar.openbioinformatics.org/) with RefGene annotation, functional prediction, snp138 annotation, in addition to SIFT and Polyphen2 predictions in column 9. The last column contains genotype counts within cases of relevant phenotypes (homozygous reference, heterozygous, homozygous alternative).

| Linkage Region | Phenotype | Chr | Start (Ref/Alt allele) | Gene | Function | snp138 | SIFT | Polyphen2 | Genotype counts |
| --- | --- | --- | --- | --- | --- | --- | --- | --- | --- |
| 11p15.4 | Depression | chr11 | 3661000 (C/T) | ART5 | exonic NS | NA | 0.05 | 0.38 | Depression: 12,4,0 |
| 11p15.4 | Depression | chr11 | 6662520 (G/A) | DCHS1 | exonic NS | rs371390544 | 0.07 | 1 | Depression: 12,4,0 |
| 12q24.22-12q24.32 | Depression | chr12 | 124171449 (G/A) | TCTN2 | exonic NS | NA | 0.38 | 0.312 | Depression: 12,4,0 |
| 1q21.1-1q21.3 | Super | chr1 | 150199027 (C/T) | ANP32E | exonic NS | NA | 1 | 0 | Super: 18,3,0 |
| 1q21.1-1q21.3 | Super | chr1 | 152080550 (G/T) | TCHH | exonic NS | NA | 0.76 | 0.411 | Super: 16,5,0 |
| 2p23.1-2p22.3 | Super | chr2 | 33817255 (G/T) | FAM98A | exonic NS | NA | 0.1 | 0.996 | Super: 14,7,0 |

## Discussion

We have identified, in rural Brazil, a family with 111 cases of mood disorder, including manic-depression. This is one of the largest mood disorder families found worldwide. We successfully performed a whole-genome linkage analysis on this large multi-generational family and after correcting for multiple testing, four regions on chromosomes 2p23.1-p22.3, 3p25.3-p24.1, 11p15.4, and 12q24.22-q24.32 achieved genome-wide significance and four regions on chromosomes 1p22.2-p21.2, 1q21.1-q21.3, 12p13.32, and 22q11.21-q12.1 achieved genome-wide suggestive linkage. Several of these regions overlap with and provide further support for previous linkage findings.

The findings from our study on chromosomes 11p15, 12p13.32-p13.31 and 12q24.22-q24.32 are specific to depression. The specificity of region 11p15 to depression is shown by extremely low LOD scores reported in the bipolar disorder only models. A similar pattern was observed for the chromosome 12 regions. While unipolar depression and BP disorder may share susceptibility loci in this family, these are loci that appear to only confer susceptibility to depression.

The highest LOD score reported in the BBF was found on chromosome 12q24.22-q24.32 (maximum LOD=4.74) under the depression phenotype model. Curtis et al. (2003) reported suggestive linkage (maximum LOD=2.8) on chromosome 12q24.31-q24.32 in seven families with multiple cases of BPD and unipolar depression. Similar to the BBF findings, they identified this region under a dominant model and including unipolar cases as affected only. Shink et al. [21] reported significant linkage in bipolar families from the Saguenay-Lac-St-Jean population of Québec. This region overlaps one identified by Ewald et al. [22], who investigated 12q22-q24 in two Danish bipolar families. The 12q23–q24 regions was also reported by Morissette et al. [23] in a large pedigree with BPD and unipolar depression from Québec.

Whole exome sequencing of 12q24.22-q24.32 showed a number of genes with mutations. One gene had more evidence with 4 depression cases carrying a mutation in the TCTN2 gene. This gene encodes a type I membrane protein that belongs to the tectonic family. Studies in mice suggest that this protein may be involved in hedgehog signaling, and essential for ciliogenesis. Mutations in this gene are associated with Meckel-Gruber Syndrome (MKS; OMIM 249000) - the most common form of syndromic neural tube defect - and Joubert syndrome (JS; OMIM 213300), which is marked by ataxia, hypotonia and other features. Both syndromes were not observed in family members but do point to the neuronal importance of TCTN2. However, this finding is tentative and requires further study within the family and replication.

We found a genome-wide significant linkage on chromosome 2p23.1-p22.3, under the super phenotype model. The region is 7.5 Mb distal to a region of possible linkage reported by [24] in Portuguese Island families with multiple patients suffering from BPD and schizophrenia. We found a convincing mutation in FAM98A that was shared by 7 of 14 cases with high read depth sequencing data and good quality scores. However, the gene encodes a protein of broadly unknown function that is part of family of proteins (including the paralogous expressed across a large of species and tissues called the DUF2465 superfamily.

The region with the most substantial evidence for linkage was chromosome 11p15.4, under the depression phenotype model. Approximately 2.6 Mb telomeric to the BBF linkage peak, Zubenko et al. [25] reported a depression susceptibility locus with significant evidence for linkage (maximum LOD=4.20) using recurrent MDD families. The findings of Zubenko et al. [25] are difficult to interpret given the thirteen chromosomal regions reaching significant linkage in their analysis. However, this region has also been implicated in bipolar disorder [26}. Our sequencing results yielded a relatively large number of plausible mutations that we were unable to select between using objective criteria. For example, mutations have been reported in the cadherin receptor DCHS1 that lead to a recessive syndrome in humans that includes periventricular neuronal heterotopia [27].

Chromosome 22q11.21-q12.1 achieved high genome-wide suggestive evidence for linkage in the BBF (LOD=3.76) under a recessive mood of disease transmission for the broad phenotype. Genome-wide significance was also previously obtained for loci on chromosome 22q12 for BPI using NPL methods and parametric methods [28]. Moreover, chromosome 22q11.2 is deleted in velocardio-facial syndrome, as first reported by Driscoll [29], which is associated with schizophrenic symptoms [30]. This locus shows a substantial decline in LOD scores observed with the inclusion of depression cases in analysis. We did not find mutations in this region that pass our QC filters.

Other regions, where we find no sequencing variants after filtering in the regions, included a suggestive linkage peak for bipolar disorder on chromosome 1p22.2-p21.3 under the narrow (maximum LOD=2.96) and broad phenotype models (maximum LOD=2.93). This region harbours the MIR137 (microRNA 137) schizophrenia locus [31]. A second region identified on 1q21.1-q21.2 is approximately 6 Mb centromeric from of a putative bipolar disorder locus (1q23.3) identified in Ashkenazi Jewish families [32], and multiplex bipolar families [33] [34]. The linkage peak on 12p13.32 had been previously implicated in both BPD and schizophrenia. This region was implicated in general mood disorders in a study of Columbian bipolar families [35]. The second most significant linkage region was found on chromosome 3p25.3-p24.1 identified using the narrow phenotype model and a dominant mode of inheritance. In related phenotypes, a larger region on chromosome 3p26-p21 was implicated in Indonesian families with schizophrenia [36] and the nearby 3p25-24 in severe recurrent depression [37].

In conclusion, we have found strong evidence for linkage in regions for bipolar disorder, broad mood disorders and also for depression alone. Some of the regions are underlain by mutations in neuronally expressed genes that are carried by affected family members. The TCTN2 and, in particular, FAM98A mutations are of interest. However, our mutation findings must be regarded as preliminary and will require replication and follow up of mutation carriers in the family. Nevertheless, this family is an unique resource, our results so far are highly promising and our follow up of these results will including integration of common variation polygenic scores as well as tracking of mutation carriers within the family.

## Acknowledgments

We gratefully acknowledge the support of the EXAC team and Daniel MacArthur for double-checking of sequence calls. **Funding**: This paper represents independent research part-funded by FAPESP, the National Institute for Health Research (NIHR) Biomedical Research Centre at South London and Maudsley NHS Foundation Trust and King’s College London. The project has received part-funding from the European Union’s Horizon 2020 research and innovation programme under Marie Sklodowska-Curie grant 658195. The views expressed are those of the authors and not necessarily those of the NHS, the NIHR or the Department of Health. **Author contributions**: GB, SA, PMcG, SDJ: analysis, GB, ACSR, MJAD, AG, SIA, SDJ: manuscript, MJAD, RAB, ACSR, SIA, GB: phenotyping and experimental design. **Competing interests**: GB has been a consultant in preclinical genomics and has received grant funding from Eli Lilly ltd.

